# TaxonMatch: taxonomic integration and tree construction from heterogeneous biological databases

**DOI:** 10.64898/2026.03.18.712418

**Authors:** Michele Leone, Valentine Rech de Laval, Harriet B. Drage, Robert M. Waterhouse, Marc Robinson-Rechavi

## Abstract

Integrating taxonomic data across heterogeneous biological databases remains a major challenge in biodiversity research due to non-standardized nomenclature, incomplete synonym annotation, and inconsistencies in taxonomic hierarchies. These issues limit interoperability between key resources such as the Global Biodiversity Information Facility (GBIF), the National Center for Biotechnology Information (NCBI), and citizen science platforms such as iNaturalist.

Here, we present TaxonMatch, a scalable and reproducible framework for taxonomic reconciliation and cross-database integration. The workflow combines string-based candidate generation using TF–IDF vectorization, supervised machine learning for match classification, and lineage-aware synonym resolution to align taxonomic entities across multiple sources. By integrating both declared and implicit equivalences, TaxonMatch resolves typographical variation, synonymy, and structural inconsistencies in taxonomic data.

The framework produces a unified taxonomic structure in which equivalent entities are reconciled while preserving source-specific identifiers, provenance information, and hierarchical relationships. We evaluate its robustness across multiple classifiers and demonstrate its effectiveness in resolving ambiguous taxonomic cases that are not handled by traditional matching approaches.

We illustrate the applicability of TaxonMatch through three use cases: the construction of a unified arthropod taxonomy integrating GBIF, NCBI, and iNaturalist data; the identification of closest extant relatives of fossil taxa with molecular information; and the integration of genomic resources with conservation data from the IUCN Red List. These applications highlight the ability of the workflow to support the integration of ecological, genomic, and paleontological datasets.

TaxonMatch provides a flexible and generalizable solution for taxonomic data integration, enabling the construction of coherent and interoperable biodiversity datasets for downstream analyses in ecology, evolution, and conservation biology.

## 1 Introduction

Taxonomy underpins our understanding of biodiversity and is central to ecological and evolutionary research (Pyle & Patterson, 2016). However, the integration of taxonomic data across databases remains challenging due to non-standardized nomenclature, heterogeneous data structures, and continuously evolving classifications (Karam *et al*., 2016; Miralles *et al*., 2020). As biodiversity datasets grow in size and complexity, these discrepancies increasingly hinder data interoperability and downstream analyses.

Major taxonomic resources provide complementary but fragmented perspectives on biodiversity. The Global Biodiversity Information Facility (GBIF) aggregates large-scale occurrence and ecological data, including fossil taxa (Flemons *et al*., 2007), whereas the National Center for Biotechnology Information (NCBI) focuses on species with molecular and genomic data (Schoch *et al*., 2020). Integrating these sources enables a more comprehensive understanding of biodiversity across ecological and evolutionary scales (Theodoridis *et al*., 2018). In addition, citizen science platforms such as iNaturalist contribute dynamic and rapidly expanding taxonomic information, often including provisional or unrecognized names.

These heterogeneous sources introduce multiple types of inconsistencies, ranging from explicit synonymy to structural discrepancies across taxonomic hierarchies. For example, identical binomial names may refer to distinct species in different databases, or a single species may be assigned to different higher-level lineages depending on the source. Such ambiguities cannot be resolved through exact string matching alone and require approaches that incorporate both lexical and hierarchical information.

Existing tools address subsets of these challenges but often operate within limited scopes (Grenié *et al*., 2023). Some focus on name standardization or synonym resolution within a single database, while others provide taxonomic backbones or name resolution services without reconciling structural inconsistencies across datasets (Hinchliff *et al*., 2015; Bennett *et al*., 2016; Rees, 2014; Specker *et al*., 2024; Rader *et al*., 2023; Flannery-Sutherland *et al*., 2022; Jones *et al*., 2023). As a result, there is a need for generalizable workflows capable of integrating heterogeneous taxonomic sources in a consistent and scalable manner.

Here, we present TaxonMatch, a workflow designed for taxonomic reconciliation and cross-database integration. TaxonMatch combines string-based candidate generation, supervised machine learning, synonym resolution, and lineage-aware integration to align taxonomic entities across multiple sources. The workflow produces a unified taxonomic structure in which equivalent entities are reconciled while preserving source-specific identifiers and hierarchical relationships.

We demonstrate the applicability of this workflow through three representative use cases: (i) the construction of a unified arthropod taxonomy integrating GBIF, NCBI, and iNaturalist data; (ii) the identification of closest extant relatives of fossil taxa with available molecular data; and (iii) the integration of genomic resources with conservation status data. These examples illustrate how a unified taxonomic framework can support the integration of ecological, genomic, and paleontological datasets.

## 2 Design rationale and principles

TaxonMatch was designed not only as a taxonomic name-matching tool, but as a reproducible framework for reconciling heterogeneous taxonomic datasets into a coherent, reusable structure. It addresses a recurring problem in biodiversity informatics: biological datasets often depend on different taxonomic backbones and therefore cannot be directly combined, even when they refer to overlapping organisms. This affects ecological observations, genomic resources, fossil records, conservation assessments, and citizen-science datasets. The design is based on four principles:

### Modularity

The architecture is divided into independent processing modules, including data acquisition, preprocessing, synonym expansion, candidate generation, supervised matching, lineage-aware validation, and tree construction. Each module can be executed separately, inspected, and adapted to different taxonomic sources or downstream applications.

### Provenance preservation

TaxonMatch does not replace source-specific identifiers with a single opaque identifier. Instead, reconciled taxonomic concepts retain links to their original GBIF, NCBI, iNaturalist, or user-provided identifiers. This allows users to trace each integrated node back to the databases from which it originated and to recover source-specific information when needed.

### Conservative reconciliation

TaxonMatch prioritizes biologically reliable integration over aggressive merging. Candidate matches are accepted only when lexical similarity, synonym information, and lineage consistency provide sufficient support. Ambiguous or weakly supported cases are preserved as unresolved rather than forced into potentially incorrect matches. This design reduces the risk of propagating erroneous taxonomic relationships into downstream analyses.

### Extensibility

Although the present study focuses on GBIF, NCBI, and iNaturalist, the workflow is source-agnostic in design. Any taxonomic dataset containing names, identifiers, ranks, and lineage information can in principle be incorporated. This makes TaxonMatch applicable beyond the arthropod-focused examples presented here, including other clades, domain-specific databases, or project-specific taxonomies.

The expected input is a set of taxonomic records containing, at minimum, canonical names, database identifiers, taxonomic ranks, and lineage paths. Optional inputs include synonym tables, accepted-name relationships, and additional metadata such as molecular-data availability or conservation status. The main outputs are a reconciled taxonomic tree, cross-database identifier mappings, synonym annotations, and unresolved ambiguity records. Together, these outputs provide both an integrated taxonomy and a transparent record of the decisions made during reconciliation.

## 3 Framework overview

TaxonMatch converts heterogeneous taxonomic sources into a reconciled, provenance-preserving taxonomic structure through a sequence of modular processing steps (Figure 1). The framework separates lexical matching, synonym handling, lineage validation, and tree construction, allowing each stage to be inspected, modified, or reused independently.

**Figure 1:**
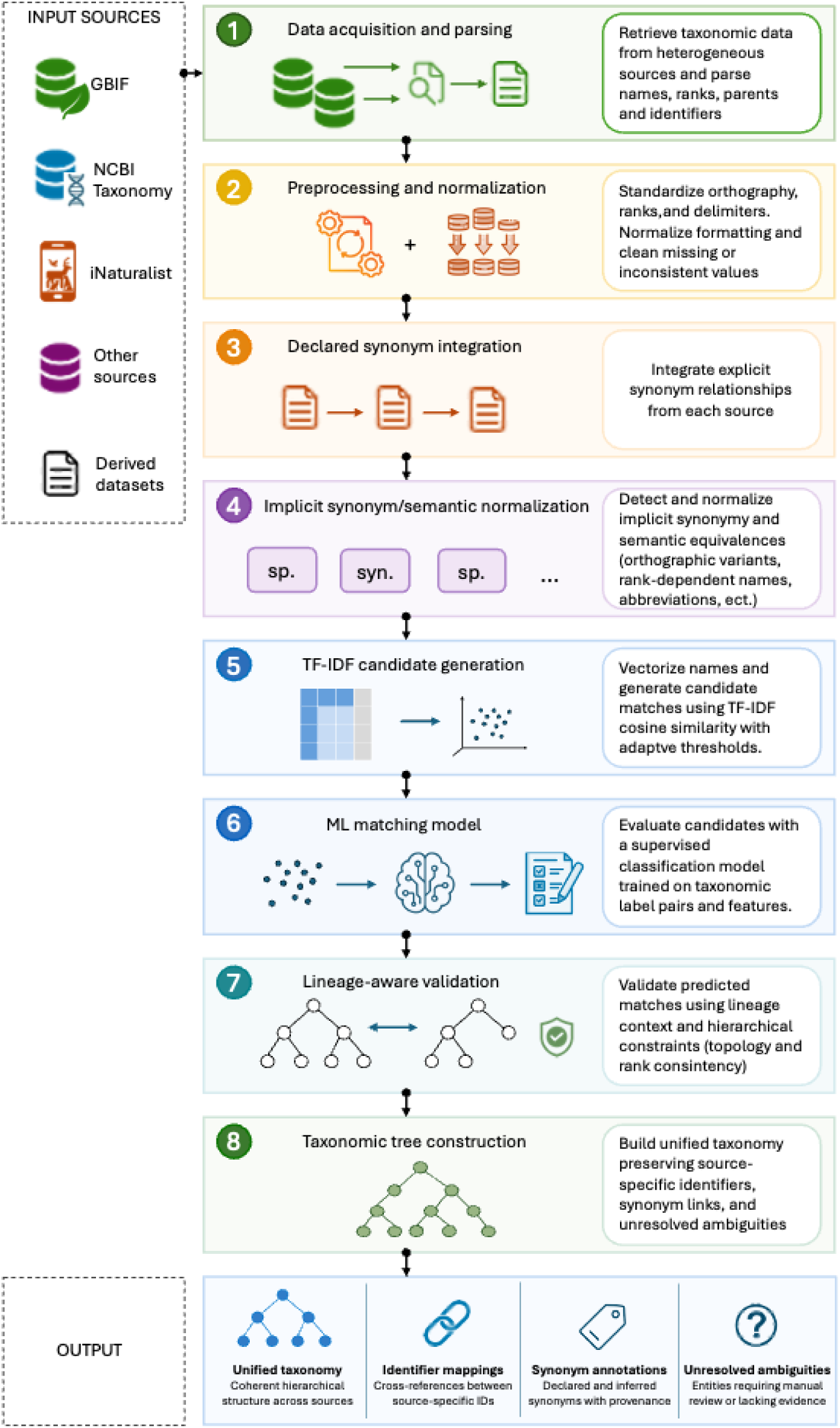
Overview of the TaxonMatch workflow. Heterogeneous taxonomic data from sources such as GBIF, NCBI, and iNaturalist are acquired, normalized, and processed through successive reconciliation modules. Declared and implicit synonym relationships are expanded through semantic normalization; candidate matches are generated using TF–IDF filtering and refined through supervised match classification. Candidate pairs are then evaluated using lineage-aware validation before being integrated into a reconciled taxonomic concept graph and unified taxonomic tree. The final outputs include a unified taxonomy, cross-database identifier mappings, synonym annotations, and unresolved ambiguity records for downstream biodiversity analyses.

The process begins with the acquisition of taxonomic datasets from structured sources such as GBIF, NCBI, and iNaturalist. For each source, TaxonMatch extracts canonical names, database identifiers, ranks, accepted-name relationships, synonym annotations, and lineage paths. These records are then normalized into a common internal representation while preserving their source-specific identifiers.

Taxonomic reconciliation is performed in three successive layers. First, declared synonym relationships are expanded into source-specific and cross-source synonym dictionaries. Second, candidate matches are generated using n-gram TF–IDF vectorization and cosine similarity, retaining only the top-ranked candidate matches for each query taxon. Third, candidate pairs are evaluated with a supervised classifier trained on string-similarity features and subsequently checked against lineage consistency constraints.

The output of these steps is not a flat list of matched names, but a reconciled taxonomic concept graph from which a unified tree can be constructed. Equivalent taxa are consolidated into shared nodes, while source-specific identifiers and synonyms are retained as annotations. Taxa that cannot be reconciled with sufficient support remain as distinct unresolved entities, preventing uncertain matches from being forced into the integrated hierarchy.

The resulting structure can be used directly as a taxonomic backbone for downstream biodiversity analyses, including cross-database querying, integration of molecular and ecological resources, fossil–extant comparisons, and conservation-genomics applications.

## 4 Reconciliation procedure

TaxonMatch consists of a sequence of deterministic and data-driven operations that transform heterogeneous taxonomic datasets into a unified and non-redundant taxonomic structure. The framework operates on multiple input sources and produces a reconciled taxonomy while preserving source-specific identifiers and hierarchical relationships. The reconciliation procedure can be summarized as the following sequence of operations:

1. **Input acquisition** Taxonomic datasets are retrieved from multiple sources, including GBIF, NCBI, and iNaturalist. Each dataset provides taxonomic entities with associated canonical names, identifiers, taxonomic ranks, and lineage information.
2. **Preprocessing and normalization** Taxonomic strings are standardized by lowercasing, removing formatting inconsistencies, and extracting lineage paths. Only accepted names are retained by default, while synonym metadata is preserved for downstream processing.
3. **Synonym expansion and semantic normalization** A synonym dictionary is constructed from declared synonym relationships in GBIF and NCBI. In addition, implicit equivalences are inferred through lineage-aware analysis, enabling the identification of semantically equivalent taxa that are not explicitly annotated as synonyms. This includes cases such as orthographic variation (e.g., *Apis mellifica* vs *Apis mellifera*) and rank-dependent representations (e.g., species vs nominotypical subspecies), which cannot be resolved through exact string matching alone.
4. **Candidate generation (TF–IDF filtering)** Each taxonomic string is transformed into an n-gram-based TF–IDF vector representation. Cosine similarity is computed between query and target datasets, and the top-*k* candidate matches are retained for each query (default: *k* = 3). This step prioritizes high recall while reducing the computational complexity of subsequent matching.
5. **Feature extraction** For each candidate pair, a feature vector is constructed using multiple string similarity metrics, including Levenshtein distance (Levenshtein, 1965), Damerau–Levenshtein distance (Damerau, 1964), Jaro–Winkler similarity (Winkler, 1990), and Hamming distance (Hamming, 1950) when applicable. These features capture both typographical variation and structural similarity between taxonomic names.
6. **Supervised matching** A trained classification model (default: XGBoost (Chen & Guestrin, 2016)) assigns a probability score to each candidate pair. A decision threshold is applied to distinguish matching from non-matching pairs. Model predictions are combined with synonym dictionaries to resolve high-confidence matches and flag ambiguous cases.
7. **Lineage-aware validation** Candidate matches are further validated using lineage consistency constraints. Pairs that exhibit strong lexical similarity but belong to incompatible higher-level taxa are rejected, while semantically equivalent taxa sharing a reconciled lineage are retained even in the absence of explicit synonym annotation. This step is critical for resolving structural inconsistencies across databases.
8. **Taxonomic integration and tree construction** Matched and unmatched entities are integrated into a unified taxonomic tree. Nodes represent taxonomic concepts linked to identifiers from multiple sources. Rule-based corrections are applied to eliminate redundant branches, resolve duplicated species-level entities, and harmonize rank-dependent representations. Synonyms are retained as annotations rather than independent nodes, ensuring a non-redundant and biologically consistent hierarchy.

The workflow is designed to be conservative: uncertain matches are not forced, and unresolved ambiguities are preserved as distinct entities. This prioritizes precision over aggressive merging and ensures that the resulting taxonomy remains interpretable and suitable for downstream biological analyses.

## 5 Input data sources

The TaxonMatch workflow integrates taxonomic data from multiple complementary biological databases, each providing distinct types of information. In this study, we focused on three primary sources: the Global Biodiversity Information Facility (GBIF), the National Center for Biotechnology Information (NCBI), and iNaturalist (iNaturalist, 2024).

GBIF provides large-scale occurrence and ecological data across a wide range of taxa, including both extant and fossil species (Global Biodiversity Information Facility, 2024). It offers a comprehensive taxonomic backbone widely used in biodiversity and ecological studies. In contrast, the NCBI taxonomy is primarily centered on species with molecular and genomic data, offering detailed lineage information and standardized identifiers used in sequence databases (Schoch *et al*., 2020). These two resources represent complementary perspectives on biodiversity: GBIF emphasizes ecological and distributional data, while NCBI focuses on molecular and genetic information.

In addition to these curated databases, we incorporated taxonomy data from iNaturalist, a citizen science platform that generates a large volume of taxonomically structured observations. While largely aligned with GBIF, the iNaturalist taxonomy includes provisional names and user-contributed annotations, providing access to taxa that may not yet be integrated into more formal taxonomic backbones.

Beyond these primary sources, the framework can be applied to datasets derived from or linked to these taxonomies. For example, the IUCN Red List relies on GBIF taxonomy for conservation assessments (International Union for Conservation of Nature, 2024), while resources such as A3Cat, Ensembl, and UniProt are structured around the NCBI taxonomy (Feron & Waterhouse, 2022; Howe *et al*., 2021; The UniProt Consortium, 2023). Similarly, the Paleobiology Database (PBDB) provides fossil records aligned with GBIF-based taxonomic structures (PBDB, 2023). By integrating these heterogeneous sources, the workflow enables the reconciliation of ecological, genomic, and paleontological data within a unified taxonomic framework.

## 6 Core components

The TaxonMatch workflow is composed of modular components that transform raw taxonomic data into a unified and consistent representation. Each component addresses a specific aspect of taxonomic reconciliation, from data acquisition to the construction of an integrated taxonomic hierarchy.

### 6.1 Data ingestion and preprocessing

Taxonomic data are automatically retrieved from external sources, including GBIF, NCBI, and iNaturalist, using dedicated scripts that download and parse the corresponding taxonomy files. From these datasets, relevant fields such as canonical names, taxonomic ranks, and lineage information are extracted and standardized.

During preprocessing, entries are filtered to retain accepted taxonomic names by default, while synonym information is preserved for downstream processing. This behavior is configurable, allowing the inclusion of provisional or non-accepted names when required for specific applications. The resulting datasets provide a normalized representation of taxonomic entities across sources.

### 6.2 Synonym resolution and semantic normalization

A central challenge in taxonomic integration lies in the resolution of synonymy and semantically equivalent entities across heterogeneous databases. While major taxonomic resources such as GBIF and NCBI provide explicit synonym annotations, a substantial proportion of equivalences remain implicit, arising from differences in nomenclatural conventions, taxonomic rank representation, and lineage structure.

TaxonMatch addresses this problem through a two-level reconciliation strategy combining explicit dictionary-based synonym mapping with lineage-aware semantic normalization.

#### Declared synonym integration

Declared synonym relationships are extracted directly from source databases. In GBIF, this includes synonym categories such as *homotypic*, *heterotypic*, and *pro parte* synonyms. In NCBI, synonym information additionally includes common names, acronyms, and alternative labels.

These relationships are aggregated into a unified synonym dictionary linking alternative names to canonical taxonomic entities. This step ensures consistent handling of formally recognized synonymy across sources.

#### Implicit synonym detection

Beyond explicitly annotated synonyms, many taxonomic inconsistencies arise from cases that are not represented as synonym relationships in the source databases. These include orthographic variation (e.g., *Apis mellifica* vs *Apis mellifera*), rank-dependent representations in which species are represented at different hierarchical levels (e.g., species vs nominotypical subspecies), and cross-database inconsistencies where equivalent taxa are assigned to different genera or higher-level lineages without explicit linkage.

Such cases cannot be resolved through exact or fuzzy string matching alone. TaxonMatch identifies these implicit equivalences by combining lexical similarity with lineage-level consistency. When two taxonomic entities share compatible higher-level ancestry and exhibit strong name similarity, they are considered candidates for semantic equivalence even in the absence of explicit synonym annotation.

For example, taxa such as *Apis mellifica mellifica* (GBIF) and *Apis mellifera mellifera* (NCBI) are not consistently linked as synonyms across databases, yet correspond to the same biological entity. Similarly, cases involving genus reassignment or orthographic variation may produce structurally distinct entries that nevertheless represent equivalent taxa.

#### Lineage-aware normalization

To resolve these ambiguities, TaxonMatch applies lineageaware normalization rules operating on both taxonomic strings and hierarchical structure. These include the alignment of canonical names across sources using shared lineage context, the normalization of redundant species-level naming in subspecies strings, the consolidation of semantically equivalent taxa into unified species-level concepts, and the preservation of infra-specific entities when explicitly represented in at least one source.

This approach ensures that equivalent biological entities are unified without collapsing distinct taxonomic concepts. Synonyms are retained as annotations associated with canonical nodes rather than treated as independent entities in the resulting taxonomy.

#### Biological consistency and conservative reconciliation

A key design principle of TaxonMatch is to prioritize biological consistency over aggressive matching. When ambiguity cannot be resolved with sufficient confidence, entities are preserved as distinct rather than forcibly aligned. This conservative strategy reduces the risk of erroneous merges while maintaining interpretability of the resulting taxonomy.

By integrating explicit synonym dictionaries with lineage-aware inference, TaxonMatch extends beyond traditional name-resolution approaches. It enables the reconciliation of taxonomic entities that are semantically equivalent but structurally divergent across databases, providing a robust foundation for downstream integration of ecological, genomic, and paleontological datasets.

### 6.3 Candidate generation

To efficiently identify potential matches between taxonomic entities, candidate generation is performed using TF–IDF vectorization of taxonomic strings. Each string is transformed into an n-gram-based TF–IDF representation capable of capturing both exact and partial lexical similarity.

For each query entity, cosine similarity is computed against entries in the target dataset, and a reduced set of top-ranked candidate matches is retained. This stage substantially reduces the search space for downstream processing while maintaining high recall of biologically plausible matches.

### 6.4 Matching model

Candidate pairs are evaluated using a supervised machine learning classifier trained to distinguish matching from non-matching taxonomic entities. The training dataset consists of balanced positive and negative pairs derived from GBIF and NCBI, including difficult non-matching cases exhibiting high lexical similarity.

Each candidate pair is represented using multiple string-similarity metrics, including Levenshtein distance (Levenshtein, 1965), Damerau–Levenshtein distance (Damerau, 1964), Jaro–Winkler similarity (Winkler, 1990), and Hamming distance (Hamming, 1950) when applicable. Together, these features capture both typographical variation and structural similarity between taxonomic names.

Among the evaluated classifiers, ensemble methods such as XGBoost demonstrated the strongest overall performance and are therefore used as the default pretrained model. The classifier assigns a prediction score to each candidate pair, which is subsequently combined with synonym dictionaries and lineage information to determine final reconciliation decisions. The machine-learning component of the workflow is shown in detail in Figure 2.

**Figure 2:**
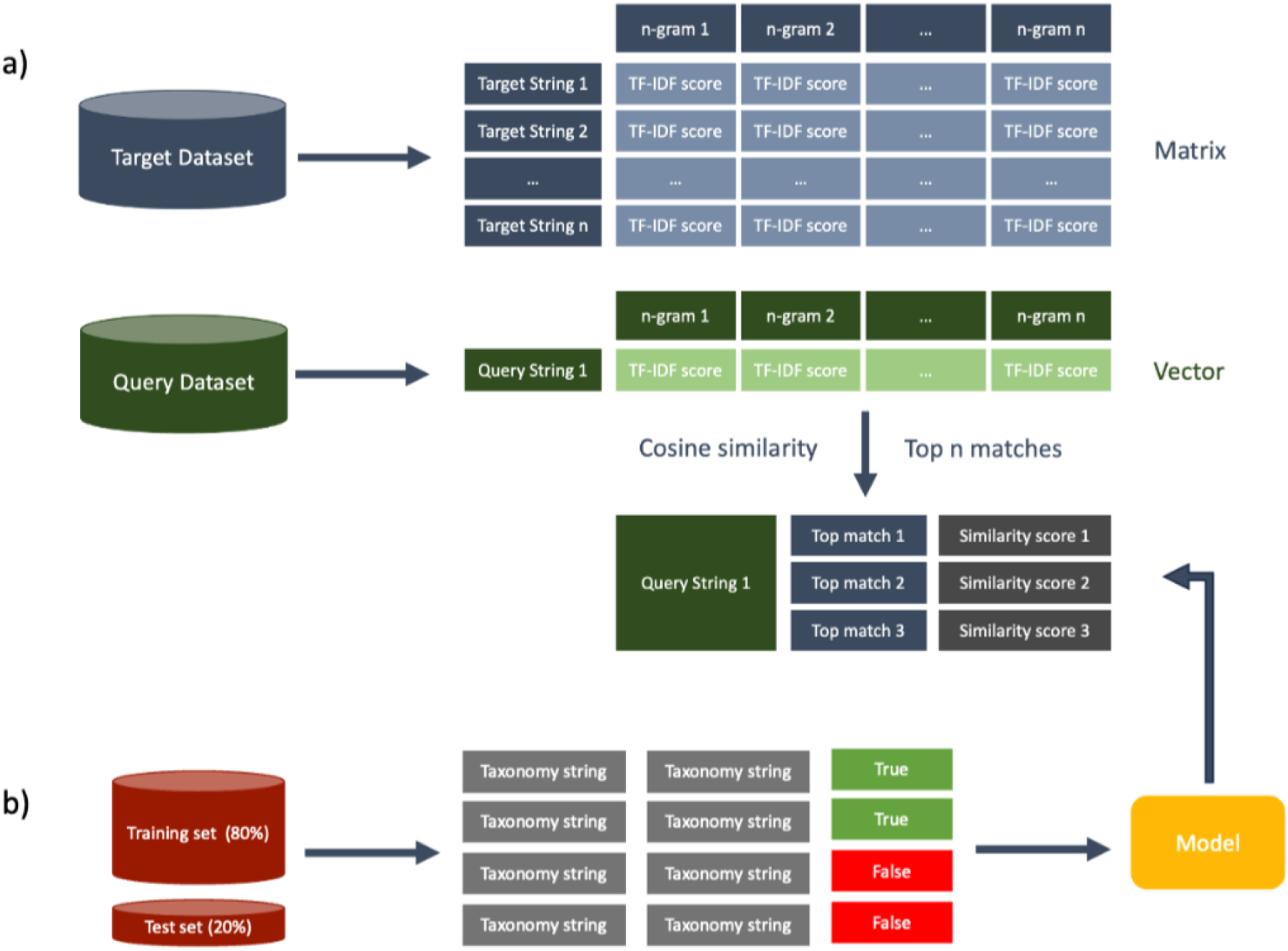
Detailed view of the machine-learning matching module used within the TaxonMatch workflow. (a) TF–IDF-based candidate generation: the target dataset is transformed into a TF– IDF matrix where each taxonomic string is represented by n-gram-based scores. Query strings are similarly vectorized, and cosine similarity is used to identify the top candidate matches for each query. (b) Model training and evaluation: training and test sets are constructed from matched and non-matched taxonomic pairs between GBIF and NCBI. A supervised classifier is trained using multiple string similarity metrics, including Levenshtein distance and Jaro– Winkler similarity, to assign a prediction score to each candidate pair.

### 6.5 Taxonomic tree construction

Matched and unmatched taxonomic entities are integrated into a unified hierarchical structure in which each node represents a taxonomic concept linked to identifiers from multiple databases.

The tree construction stage reconciles discrepancies in nomenclature and taxonomic rank while preserving source-specific information. Rule-based corrections are applied to eliminate redundant branches, resolve duplicated species-level entities, and harmonize inconsistent hierarchical representations arising from divergent taxonomic conventions.

The resulting taxonomic structure provides a coherent and non-redundant framework suitable for downstream integration of ecological, genomic, conservation, and paleontological datasets.

## 7 Error propagation analysis

The TaxonMatch workflow consists of multiple sequential processing steps, each of which may introduce specific sources of error that can influence downstream stages. Understanding how these errors propagate across the workflow is essential for evaluating its robustness and reliability.

During candidate generation, TF–IDF vectorization combined with cosine similarity is used to identify a reduced set of potential matches. This step prioritizes computational efficiency and high recall; however, if the correct match is not included among the top candidate pairs, it will not be evaluated in subsequent steps. Such cases result in missed alignments between taxonomic entities across datasets.

In the matching stage, the supervised classification model introduces both false positives and false negatives. False positives correspond to incorrect alignments between distinct taxonomic entities, while false negatives result in missed matches. These errors are partially mitigated by combining model predictions with synonym dictionaries and lineage-based validation, which provide additional constraints on candidate selection.

Errors may also arise during synonym resolution. While declared synonyms are systematically integrated from source databases, implicit or unannotated equivalences may not always be detected. In such cases, semantically equivalent taxa may remain unresolved, particularly when inconsistencies arise from differences in taxonomic rank representation or naming conventions across databases.

Importantly, the taxonomic tree construction step incorporates rule-based controls designed to prevent inconsistent or biologically implausible structures. As a result, the workflow favors conservative integration: uncertain matches are not forced, and potentially incorrect alignments are avoided. Rather than introducing erroneous merges, unresolved ambiguities are preserved as separate entities.

Overall, error propagation within the workflow primarily manifests as missed or unresolved alignments rather than incorrect structural integration. This reflects a deliberate design choice prioritizing precision and interpretability over aggressive matching, ensuring that the resulting taxonomic framework remains reliable for downstream analyses.

## 8 Model robustness analysis

To evaluate the robustness of the reconciliation procedure, we assessed classifier behaviour across multiple supervised learning approaches trained on the same taxonomic matching task. This comparison was intended to evaluate the stability of the matching stage with respect to model choice rather than to optimize predictive performance alone.

Various models were evaluated for taxonomic entity classification, including Random-Forest, XGBoost, GradientBoosting, K-Nearest Neighbors, DecisionTree, AdaBoost, Support Vector Machine, Multilayer Perceptron, and Perceptron. Performance was assessed using a training dataset comprising 50,000 samples, evenly divided between positive and negative taxonomic matches.

All models were evaluated using 5-fold cross-validation to estimate generalization performance and reduce the risk of overfitting. Among the evaluated classifiers, XGBoost demonstrated the strongest overall performance, achieving an accuracy of 0.97 together with high precision, recall, and F1 score values (Table 1). Ensemble-based approaches, particularly XGBoost and RandomForest, consistently outperformed linear and single-estimator methods across evaluation metrics.

**Table 1:**
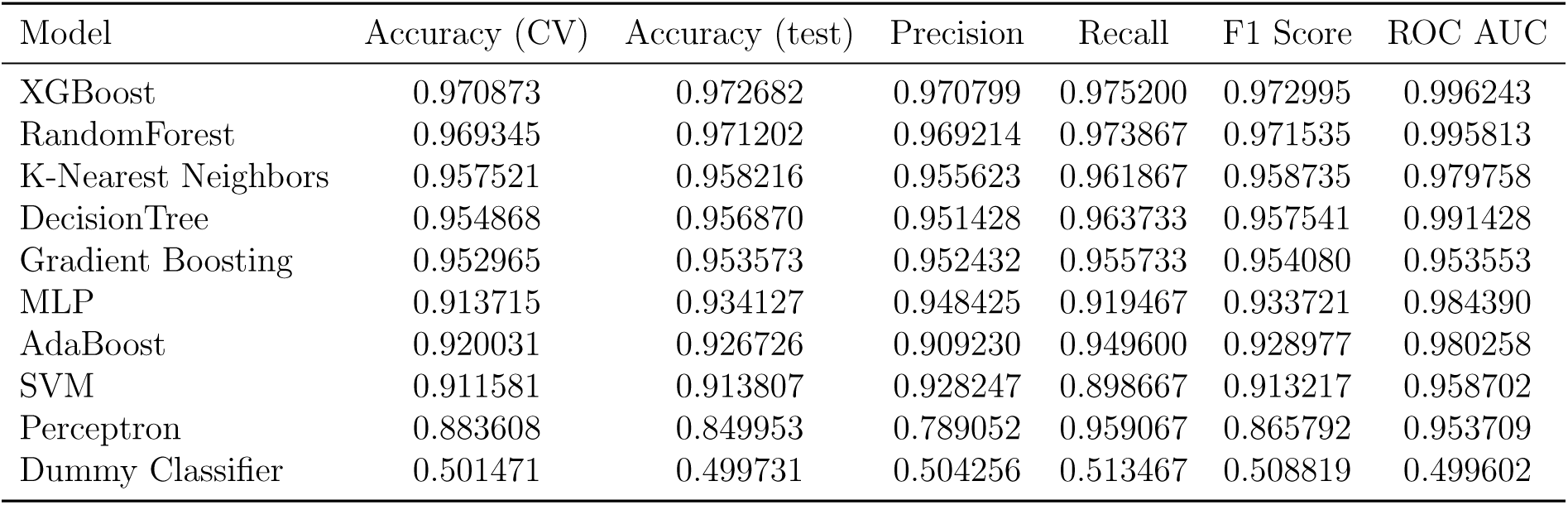
Performance comparison of classification models used in the TaxonMatch workflow. Models are ranked by cross-validation accuracy.

| Model | Accuracy (CV) | Accuracy (test) | Precision | Recall | F1 Score | ROC AUC |
| --- | --- | --- | --- | --- | --- | --- |
| XGBoost | 0.970873 | 0.972682 | 0.970799 | 0.975200 | 0.972995 | 0.996243 |
| RandomForest | 0.969345 | 0.971202 | 0.969214 | 0.973867 | 0.971535 | 0.995813 |
| K-Nearest Neighbors | 0.957521 | 0.958216 | 0.955623 | 0.961867 | 0.958735 | 0.979758 |
| DecisionTree | 0.954868 | 0.956870 | 0.951428 | 0.963733 | 0.957541 | 0.991428 |
| Gradient Boosting | 0.952965 | 0.953573 | 0.952432 | 0.955733 | 0.954080 | 0.953553 |
| MLP | 0.913715 | 0.934127 | 0.948425 | 0.919467 | 0.933721 | 0.984390 |
| AdaBoost | 0.920031 | 0.926726 | 0.909230 | 0.949600 | 0.928977 | 0.980258 |
| SVM | 0.911581 | 0.913807 | 0.928247 | 0.898667 | 0.913217 | 0.958702 |
| Perceptron | 0.883608 | 0.849953 | 0.789052 | 0.959067 | 0.865792 | 0.953709 |
| Dummy Classifier | 0.501471 | 0.499731 | 0.504256 | 0.513467 | 0.508819 | 0.499602 |

To further characterize classification behaviour, Receiver Operating Characteristic (ROC) curves were generated for all evaluated models (Supplementary Figure S1). XG-Boost and RandomForest achieved the highest Area Under the Curve (AUC) values, indicating strong discrimination between matching and non-matching taxonomic pairs even in cases involving high lexical similarity.

Overall, these results indicate that the reconciliation process remains stable across different classification strategies, while also supporting the use of XGBoost as a robust default model for large-scale taxonomic integration.

## 9 Benchmarking and validation

To evaluate the effectiveness of the TaxonMatch workflow, we performed a series of benchmarking analyses comparing its performance with existing approaches and assessing its behaviour on real-world taxonomic datasets.

First, we compared TaxonMatch with existing taxonomic integration and name-resolution resources, including the GBIF Backbone Taxonomy (Global Biodiversity Information Facility, 2024), Catalogue of Life (Bánki *et al*., 2025), Open Tree of Life (Hinchliff *et al*., 2015), Global Names Resolver and Global Names Recognition and Discovery (GNR/GNRD) (Bennett *et al*., 2016), Taxamatch (Rees, 2014), Treemendous (Specker *et al*., 2024), PhyloMatcher (Rader *et al*., 2023), and Fossilbrush together with the Palaeoverse function *tax check()* (Flannery-Sutherland *et al*., 2022; Jones *et al*., 2023).

These resources address different aspects of taxonomic reconciliation, including curated backbone construction, name resolution, fuzzy matching, synonym handling, and taxonomic anomaly detection. However, they generally focus either on lexical normalization, curated synonym handling, or predefined consensus hierarchies, whereas TaxonMatch combines candidate generation, supervised classification, synonym integration, and lineage-aware validation within a single reconciliation framework.

This comparison highlights the ability of TaxonMatch to resolve both typographical variation and structural mismatches between databases. In particular, TaxonMatch is able to identify valid matches in cases where traditional approaches fail due to strict string matching constraints or incomplete synonym coverage.

To further evaluate the behaviour of the workflow, we analysed ambiguous taxonomic cases that are not trivially resolved through exact matching. These include pairs of taxa with high lexical similarity but different biological meanings, as well as cases where multiple candidate matches exist.

These examples demonstrate that the workflow is capable of distinguishing between true and false matches by combining multiple sources of information, including string similarity, synonym relationships, and lineage consistency.

We also compared the resulting taxonomic structure with the Open Tree of Life (OTOL) framework (Hinchliff *et al*., 2015), a widely used reference for unified taxonomic integration. While OTOL focuses on integrating phylogenetic and taxonomic information, TaxonMatch provides a complementary approach based on data-driven reconciliation across heterogeneous sources. The comparison highlights differences in coverage and alignment strategies, with TaxonMatch offering improved integration of datasets that are not directly aligned in existing taxonomic backbones.

**Table 2:**
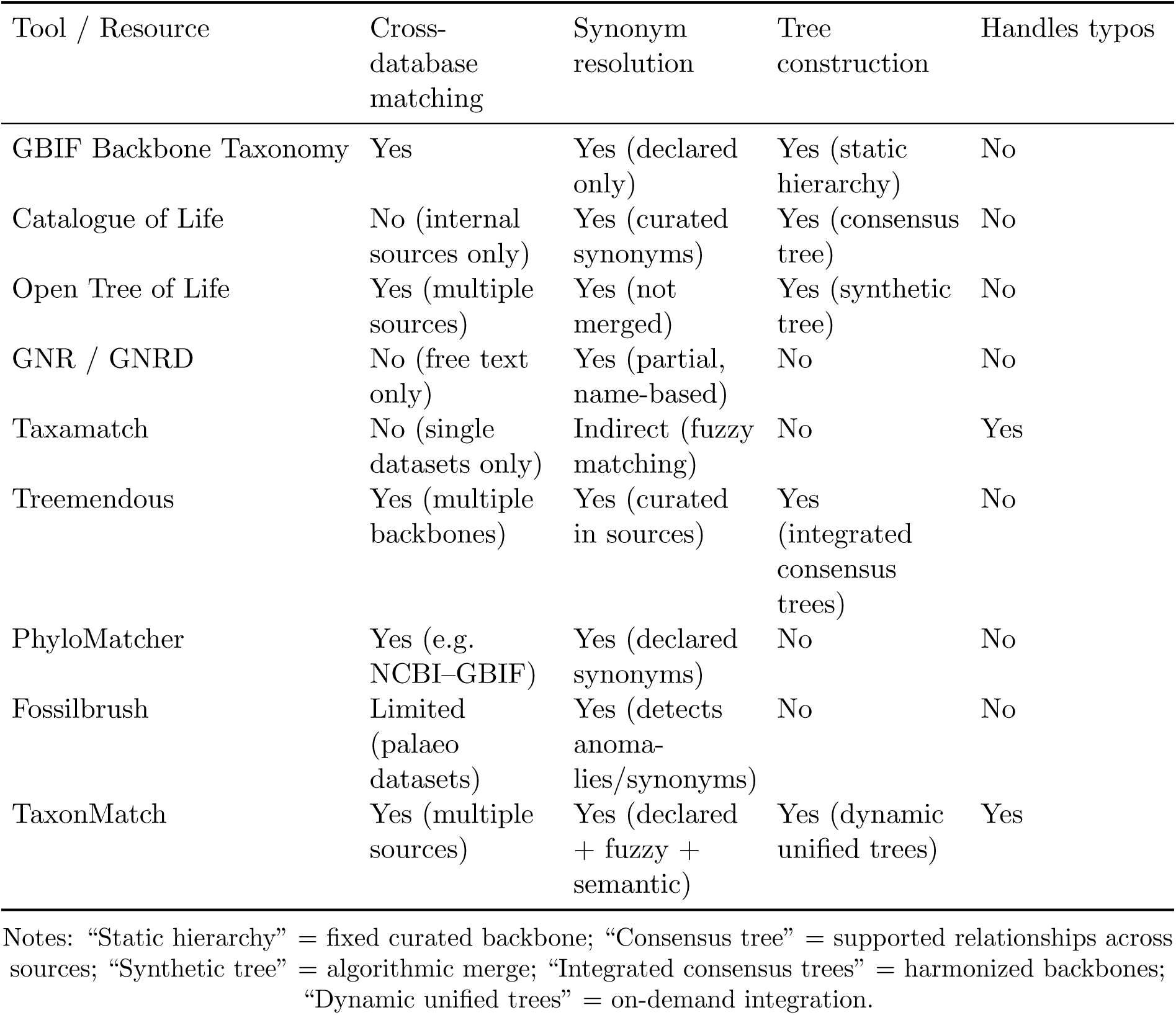
Comparison of major taxonomic integration tools and services.

| Tool / Resource | Cross-database matching | Synonym resolution | Tree construction | Handles typos |
| --- | --- | --- | --- | --- |
| GBIF Backbone Taxonomy | Yes | Yes (declared only) | Yes (static hierarchy) | No |
| Catalogue of Life | No (internal sources only) | Yes (curated synonyms) | Yes (consensus tree) | No |
| Open Tree of Life | Yes (multiple sources) | Yes (not merged) | Yes (synthetic tree) | No |
| GNR / GNRD | No (free text only) | Yes (partial, name-based) | No | No |
| Taxamatch | No (single datasets only) | Indirect (fuzzy matching) | No | Yes |
| Treemendous | Yes (multiple backbones) | Yes (curated in sources) | Yes (integrated consensus trees) | No |
| PhyloMatcher | Yes (e.g. NCBI-GBIF) | Yes (declared synonyms) | No | No |
| Fossilbrush | Limited (palaeo datasets) | Yes (detects anomalies/synonyms) | No | No |
| TaxonMatch | Yes (multiple sources) | Yes (declared + fuzzy + semantic) | Yes (dynamic unified trees) | Yes |

Notes: “Static hierarchy” = fixed curated backbone; “Consensus tree” = supported relationships across sources; “Synthetic tree” = algorithmic merge; “Integrated consensus trees” = harmonized backbones; “Dynamic unified trees” = on-demand integration.

Overall, these benchmarking analyses demonstrate that the TaxonMatch workflow provides a robust and flexible solution for taxonomic reconciliation, outperforming traditional string-based approaches and enabling the integration of diverse biological datasets.

**Table 3:**
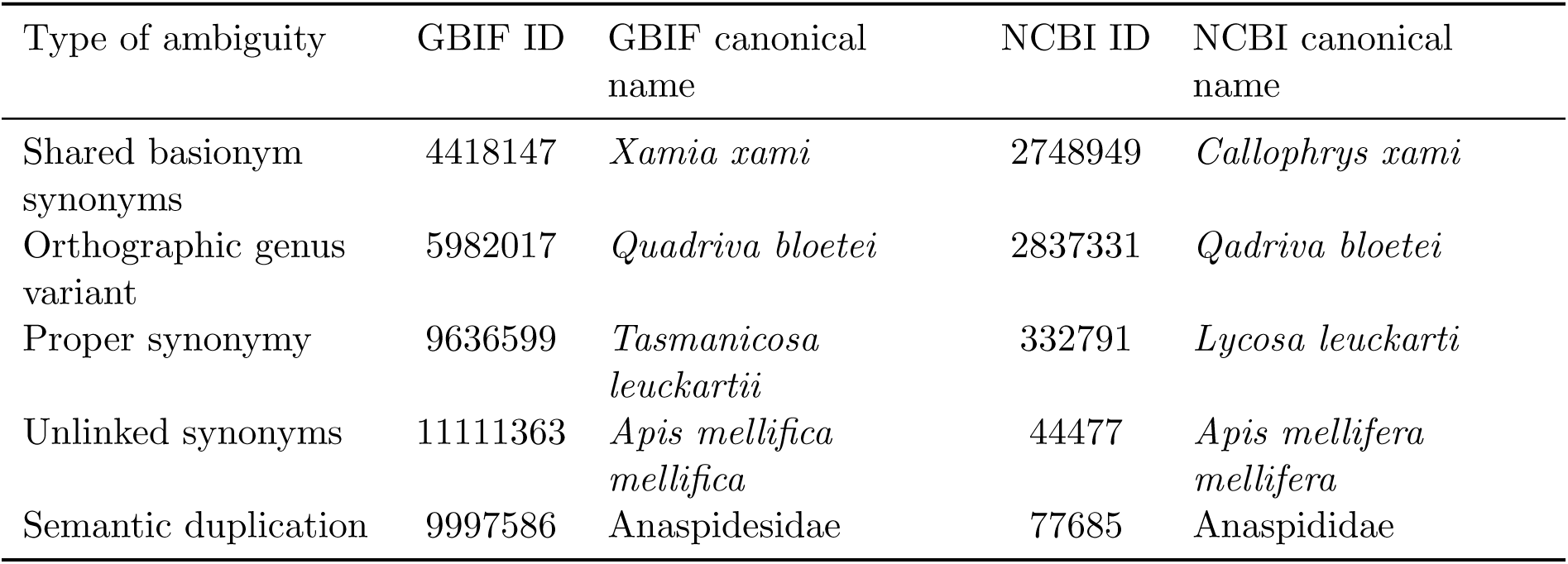
Arthropod examples of unresolved taxonomic ambiguities in GBIF and NCBI that are correctly interpreted by TaxonMatch. These cases involve semantically equivalent taxa that are structurally divergent across databases due to inconsistent naming conventions, typographical variations, or invalid hierarchical assignments. They are harmonized by TaxonMatch through lineage correction and semantic normalization.

**Table 4:** Examples of non-declared taxonomic matches inferred by TaxonMatch in the absence of explicit synonym or cross-reference annotations, identified through lineage-aware reconciliation.

| GBIF taxon name | GBIF taxon ID | NCBI taxon name | NCBI taxon ID |
| --- | --- | --- | --- |
| Iolanthe rostrata | 9981640 | Acanthaspidia rostratus | 263151 |
| Atychia funebris | 5141599 | Brachodes funebris | 2743946 |
| Pachylomera femoralis | 4996751 | Pachylomerus femoralis | 205304 |
| Hygrochroa albipunctata | 1868598 | Apatelodes albipunctata | 1093843 |
| Platphalonia mystica | 12244282 | Platphalonidia mystica | 1558185 |

## 10 Applications

We demonstrate the applicability of the TaxonMatch workflow through three representative use cases involving the integration of heterogeneous biodiversity datasets across ecological, genomic, and paleontological domains.

### 10.1 Backbone taxonomy construction

As a first application, we constructed a unified arthropod taxonomy by integrating data from GBIF (Global Biodiversity Information Facility, 2024), NCBI (Schoch *et al*., 2020), and iNaturalist (iNaturalist, 2024). This backbone enables the integration of ecological observations, genomic data, and citizen-science records within a single taxonomic framework, as required by resources such as MoultDB.

Across these sources, only a limited fraction of taxa are shared, while the majority remain exclusive to individual datasets, highlighting the extent of fragmentation in current taxonomic resources. The overlap between the three datasets is illustrated in Figure 3, showing that most taxa are unique to a single source, with only a small proportion shared across all platforms.

**Figure 3:**
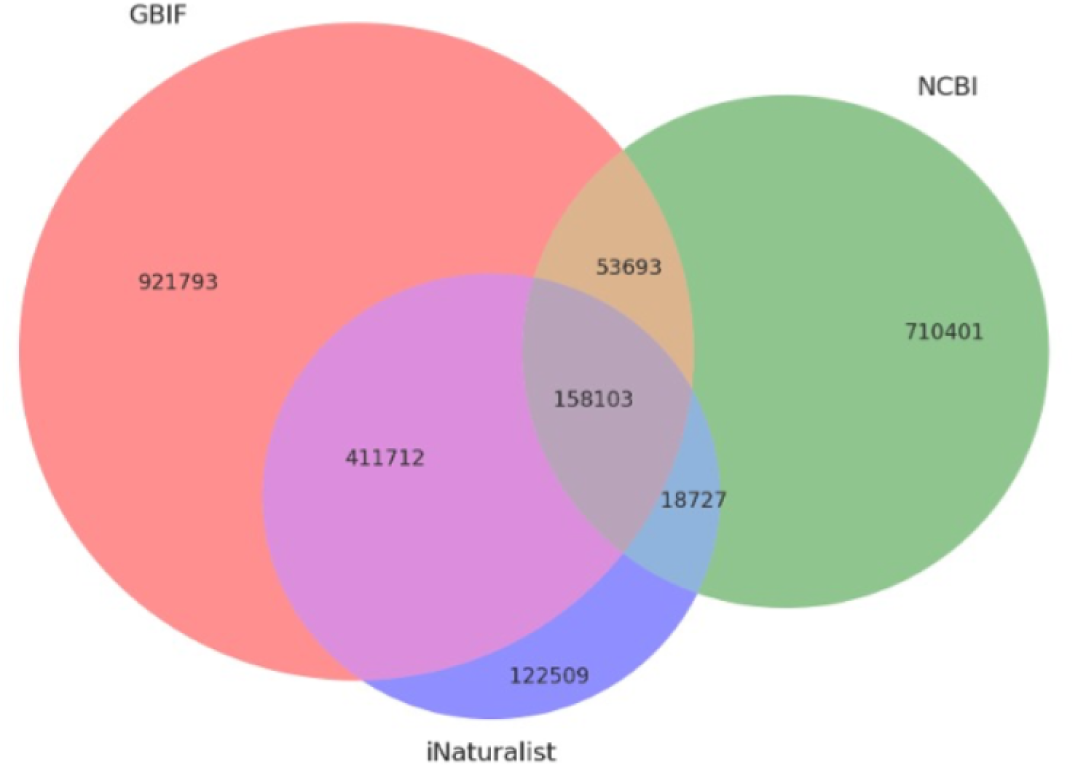
Venn diagram showing the overlap of arthropod taxonomic entries across GBIF, NCBI, and iNaturalist.

The integrated backbone taxonomy produced by TaxonMatch contains 2,396,938 unique taxonomic identifiers, corresponding to 1,458,614 species-level entities. This demonstrates the scalability of the workflow when applied to large and heterogeneous datasets.

By resolving synonymy, correcting structural inconsistencies, and preserving source-specific identifiers, the workflow produces a coherent and non-redundant taxonomy. The resulting structure supports cross-database queries and downstream analyses that would not be possible using individual taxonomic sources in isolation.

An example of the resulting integrated taxonomy is shown in Figure 4, which illustrates the reconciliation of identifiers from GBIF, NCBI, and iNaturalist within a single taxonomic tree.

**Figure 4:**
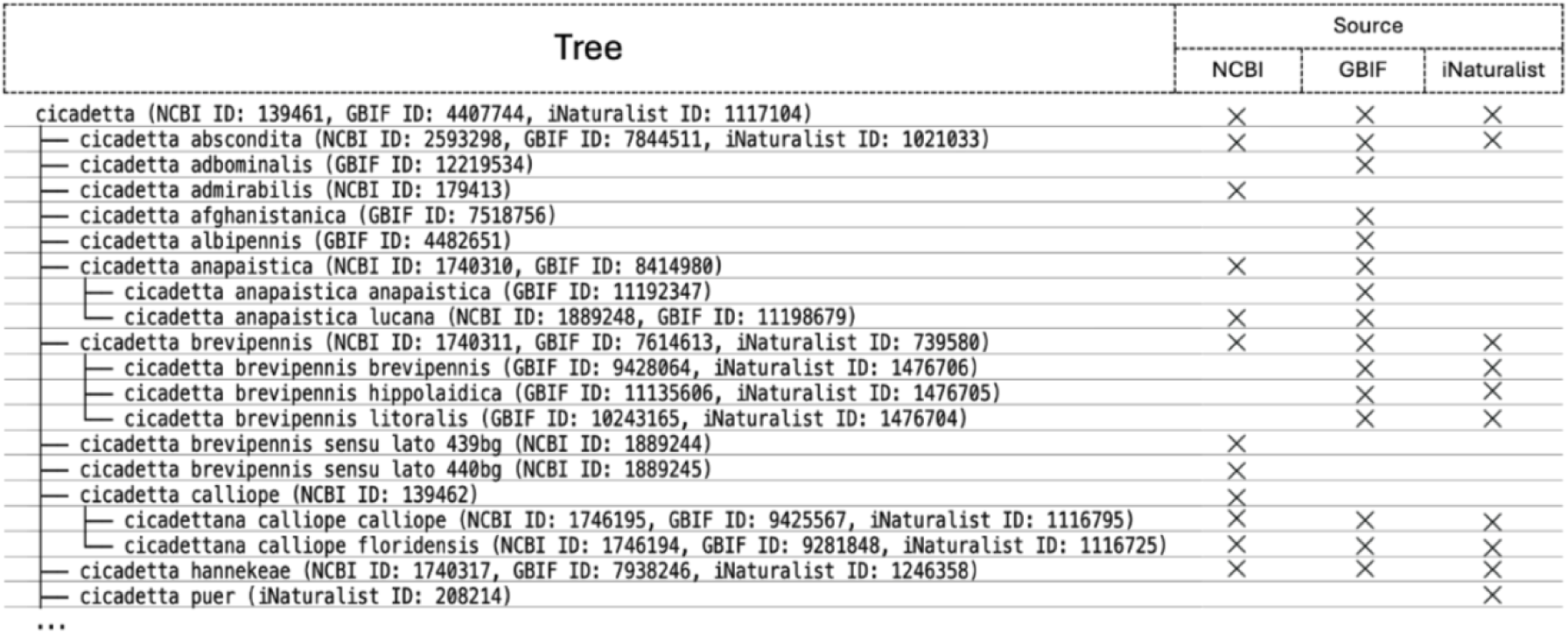
Taxonomic tree of the genus *Cicadetta*, showing integrated identifiers from NCBI, GBIF, and iNaturalist. The reconciled tree includes taxa present in only one source as well as shared entities across databases.

### 10.2 Fossil–extant mapping

As a second application, we used TaxonMatch to identify the closest extant relatives of fossil taxa with available molecular data. This use case demonstrates how the workflow enables integration between paleontological and genomic datasets.

We focused on the extinct crustacean *Ristoria pliocaenica*, combining taxonomic information from GBIF-derived sources with molecular data from NCBI. Using lineage-aware matching, the workflow identifies the closest shared clade between the fossil taxon and extant species, and reconstructs the corresponding taxonomic subtree.

A portion of the reconstructed taxonomy is shown in Figure 5, highlighting extant relatives within the family Leucosiidae. For these taxa, molecular data are available at the marker level, although complete genome assemblies are not present.

**Figure 5:**
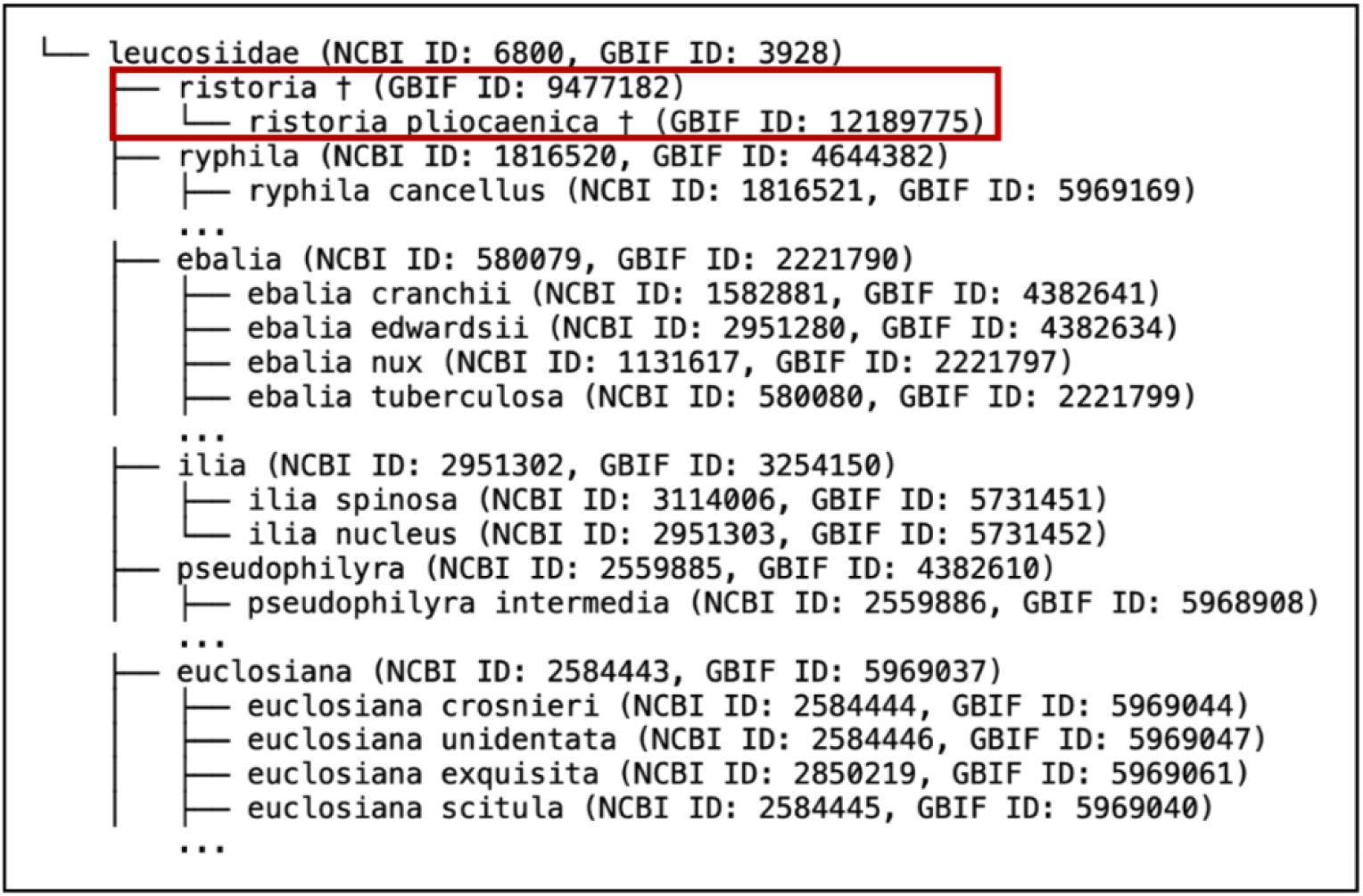
Reconstructed taxonomic subtree centered on *Ristoria pliocaenica*, with extant relatives in the family Leucosiidae. The tree is based on taxonomic alignment between GBIF-derived sources and NCBI. *Ristoria pliocaenica* is highlighted, while living taxa are shown without emphasis. Each node displays the scientific name, NCBI Taxonomy ID, and GBIF ID.

This approach enables evolutionary analyses across deep time by linking fossil taxa to extant species with available molecular data. More generally, the workflow can identify relevant comparison groups even when direct species-level matches are not available, by leveraging shared lineage structure.

### 10.3 Genomics and conservation integration

In a third application, we used TaxonMatch to integrate genomic resources with conservation data by aligning species from the Arthropoda Assembly Assessment Catalogue (A3Cat) with the IUCN Red List.

Because A3Cat (Feron & Waterhouse, 2022) is based on NCBI taxonomy and the IUCN Red List (International Union for Conservation of Nature, 2024) follows GBIF taxonomy, this integration requires cross-database reconciliation. TaxonMatch enables the alignment of these resources, allowing the identification of species for which both genome assemblies and conservation status are available.

We identified 177 arthropod species with both genomic data and IUCN classification. Among these, several taxa are classified as Vulnerable, Endangered, or Critically Endangered, indicating that genomic resources remain limited for species of highest conservation concern. The distribution of species with available genome assemblies across conservation categories is shown in Supplementary Figure S2. A subset of these species is reported in Table 5.

**Table 5:**
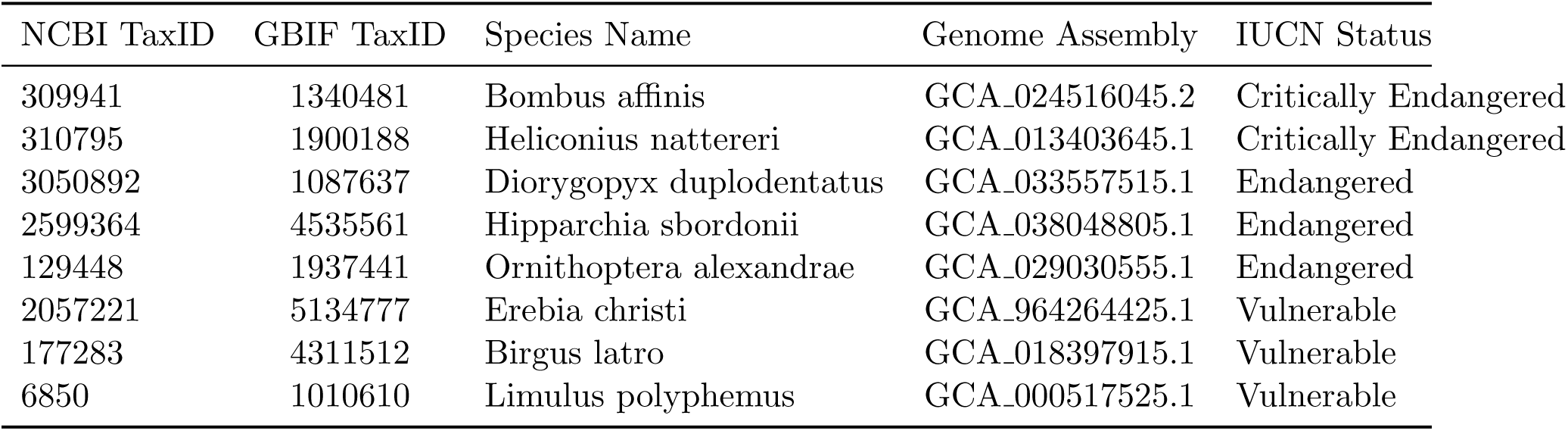
Examples of arthropod species for which TaxonMatch successfully aligned genomic records from A3Cat with conservation statuses from the IUCN Red List.

Beyond these examples, the workflow enables the systematic identification of species lacking genomic resources across conservation categories (Supplementary Figure S3). This highlights gaps in current sequencing efforts and provides a framework for prioritizing species in conservation genomics.

Overall, these applications illustrate how the TaxonMatch workflow supports the integration of large-scale and heterogeneous biodiversity datasets, enabling analyses that combine ecological, genomic, and evolutionary information within a unified taxonomic framework.

## 11 Discussion

The main contribution of TaxonMatch lies not in individual algorithmic components alone, but in their integration into a reproducible reconciliation framework for heterogeneous taxonomic resources. Rather than functioning solely as a name-matching system, TaxonMatch combines candidate generation, supervised classification, synonym integration, and lineage-aware validation into a unified procedure for constructing interoperable taxonomic structures across biodiversity databases.

This distinction is important because taxonomic reconciliation is increasingly becoming an infrastructure problem rather than a purely nomenclatural one. Ecological observations, genomic repositories, conservation datasets, fossil records, and citizen-science resources frequently depend on partially incompatible taxonomic backbones, limiting interoperability between biological data sources. By preserving provenance information while reconciling semantically equivalent entities, TaxonMatch enables the construction of integrated taxonomic representations suitable for downstream biodiversity analyses.

A key strength of TaxonMatch is its ability to resolve not only declared synonymy, but also implicit and structural inconsistencies that arise from differences in taxonomic representation across databases. These include orthographic variation, rank-dependent naming, and semantically equivalent taxa distributed across divergent taxonomic hierarchies. Such cases are not captured by synonym dictionaries alone and cannot be reliably resolved through exact or fuzzy string matching. By integrating lexical similarity with lineage-level constraints, TaxonMatch enables the reconciliation of taxonomic entities that are structurally distinct but biologically equivalent.

Another important aspect of the framework is its conservative design. Rather than forcing alignment between uncertain candidates, TaxonMatch prioritizes precision by combining TF–IDF-based candidate filtering, supervised classification, and lineage-aware validation. This reduces the risk of erroneous merges, which can introduce artificial relationships and propagate errors in downstream phylogenetic or comparative analyses. When ambiguity cannot be resolved with sufficient confidence, entities are preserved as distinct, maintaining interpretability and biological validity.

The applications presented in this study illustrate the versatility of this reconciliation strategy. The construction of a unified arthropod taxonomy demonstrates its scalability and its capacity to integrate large and heterogeneous datasets. The fossil–extant mapping highlights its relevance for evolutionary analyses across deep time, while the integration of genomic and conservation data shows its potential for supporting conservation genomics and biodiversity prioritization. These use cases emphasize that taxonomic reconciliation is not an isolated task, but a prerequisite for integrating ecological, genomic, and paleontological data.

Compared to existing tools, TaxonMatch provides a flexible and extensible framework that operates directly on structured taxonomies rather than on isolated name strings or fixed backbones. While resources such as the GBIF Backbone Taxonomy and the Open Tree of Life provide valuable reference structures, they rely on predefined hierarchies and do not explicitly address cross-database inconsistencies or implicit equivalences. Similarly, name-resolution services and fuzzy-matching approaches are effective for lexical normalization but do not incorporate lineage-level reasoning. TaxonMatch complements these approaches by enabling data-driven reconciliation across heterogeneous sources, preserving both hierarchical structure and source-specific identifiers.

Despite these strengths, several limitations remain. The performance of the matching model depends on the availability and representativeness of training data, and may be affected by biases in the underlying taxonomic sources. In addition, while lineage-aware validation improves robustness, it cannot fully resolve cases where taxonomic hierarchies are themselves inconsistent or incomplete. The current implementation focuses primarily on taxonomies derived from GBIF and NCBI, and its behaviour when applied to entirely independent or conflicting taxonomic systems remains to be evaluated.

Future developments could extend TaxonMatch by incorporating additional taxonomic sources, integrating phylogenetic information directly into the reconciliation process, and introducing explicit confidence estimates for matched entities. Such extensions would further strengthen its role as a general framework for taxonomic integration.

Overall, TaxonMatch provides a scalable and biologically grounded framework for taxonomic reconciliation, enabling the construction of coherent and interoperable taxonomic structures that support a wide range of applications in ecology, evolution, and conservation biology.

## Author contributions

M.L. and V.R.d.L. conceived the project. M.L. designed and implemented the method, developed the software, and performed the analyses. V.R.d.L. contributed to the use and validation of taxonomic data. R.M.W. provided input on genomic and biodiversity applications. H.B.D. provided input on paleontological applications. M.R.R. supervised the project. M.L. wrote the manuscript with input from all authors.

## Acknowledgements

This work was supported by a Sinergia grant (no. CRSII5 198691) from the Swiss National Science Foundation. We thank all members of the Sinergia project for valuable discussions.

## Data and code availability

All data and source code used in this study are publicly available at: https://github.com/MoultDB/TaxonMatch.

The repository includes the codebase, example datasets, and instructions required to reproduce the analyses presented in this study.

