## Supplementary for "TaxonMatch: taxonomic integration and tree construction from heterogeneous biological databases"

### Supplementary Material
### TaxonMatch

#### Supplementary Figure S1. ROC curves of all classifiers

Receiver Operating Characteristic (ROC) curves for all evaluated classifiers, including ensemble methods (XGBoost, Random Forest, Gradient Boosting, AdaBoost), linear models (Perceptron and Support Vector Machine), neural networks (Multilayer Perceptron), instance-based methods (K-Nearest Neighbors), decision trees, and a baseline Dummy Classifier.

The curves illustrate the trade-off between true positive rate (TPR) and false positive rate (FPR) across classification thresholds. Ensemble methods, particularly XGBoost and RandomForest, achieved the highest AUC values, indicating strong discriminatory power for distinguishing matching from non-matching taxonomic entities.

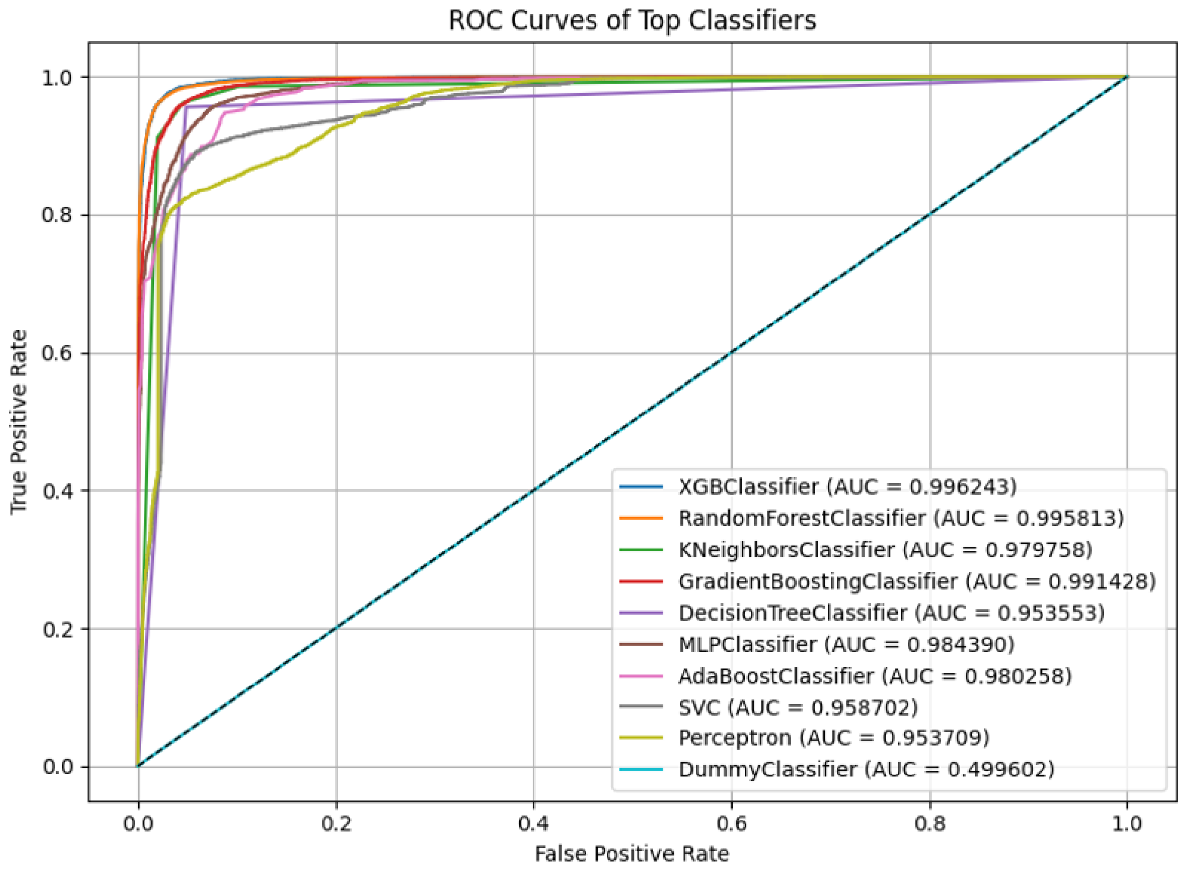


Supplementary Figure S1

### Supplementary Figure S2. Arthropod species with genome assemblies by conservation status

Distribution of arthropod species with available genome assemblies grouped according to IUCN conservation categories.

Among the 177 reconciled species identified through TaxonMatch, most were classified as Least Concern, whereas comparatively few belonged to threatened categories such as Vulnerable, Endangered, or Critically Endangered.

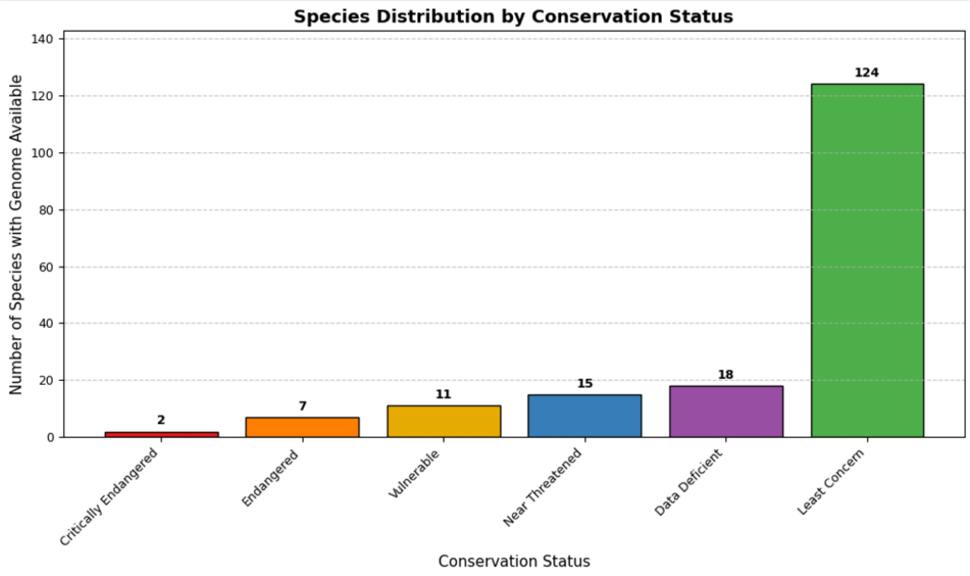


Supplementary Figure S2

### Supplementary Figure S3. Arthropod species without genome assemblies by conservation status

Distribution of arthropod species lacking genome assemblies across IUCN conservation categories.

The large discrepancy between species with and without available genomic resources highlights substantial gaps in current sequencing efforts, particularly among threatened taxa.

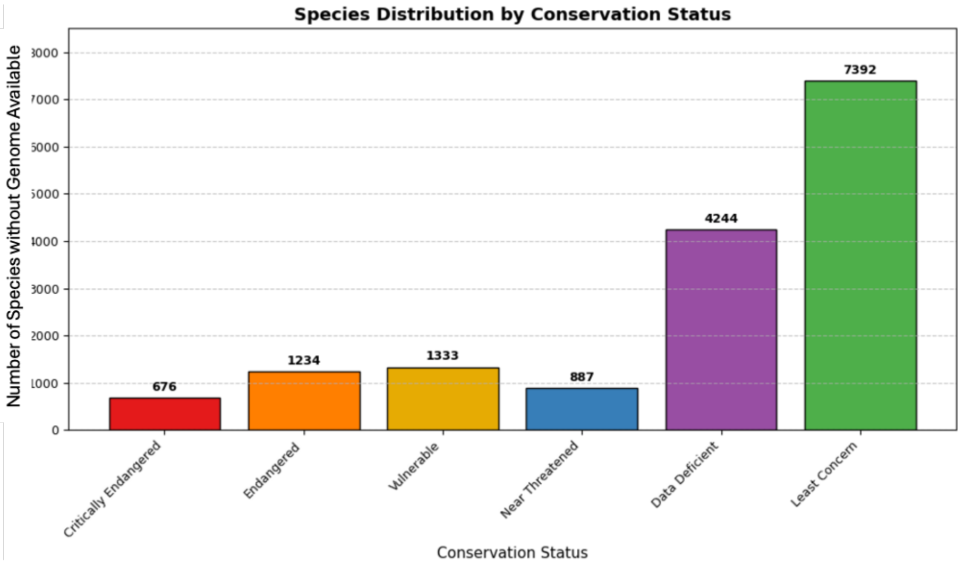


Supplementary Figure S3
